# Nested Active Learning for Efficient Model Contextualization and Parameterization: Pathway to generating simulated populations using multi-scale computational models

**DOI:** 10.1101/644401

**Authors:** Chase Cockrell, Jonathan Ozik, Nick Collier, Gary An

**Affiliations:** Department of Surgery, University of Vermont; Argonne National Laboratory

## Abstract

There is increasing interest in the use of mechanism-based multi-scale computational models (such as agent-based models) to generate simulated clinical populations in order to discover and evaluate potential diagnostic and therapeutic modalities. The description of the environment in which a biomedical simulation operates (model context) and parameterization of internal model rules (model content) requires the optimization of a large number of free-parameters. In this work, we utilize a nested active-learning workflow to efficiently parameterize and contextualize an agent-based model (ABM) of systemic inflammation used to examine sepsis.

**Methods:** Contextual parameter space was examined using four parameters external to the model’s rule-set. The model’s internal parameterization, which represents gene expression and associated cellular behaviors, was explored through the augmentation or inhibition of signaling pathways for 12 signaling mediators associated with inflammation and wound healing. We have implemented a nested active learning approach in which the clinically relevant model environment space for a given internal model parameterization is mapped using a small Artificial Neural Network (ANN). The outer AL level workflow is a larger ANN which uses active learning to efficiently regress the volume and centroid location of the CR space given by a single internal parameterization.

**Results:** We have reduced the number of simulations required to efficiently map the clinically relevant parameter space of this model by approximately 99%. Additionally, we have shown that more complex models with a larger number of variables may expect further improvements in efficiency.

## Introduction

There is increasing interest in the use of mechanism-based multi-scale computational models as an aid to more traditional biomedical research methods. This approach integrates existing cellular and molecular mechanistic knowledge into tissue-, organ- and patient-level representations that are used to generate “virtual populations” to investigate potential diagnostic and therapeutic modalities. This is the underlying concept of creating “digital twins” for the study of precision medicine (1) and the regulatory interest in advancing the ability to perform *in silico* clinical trials (2). However, achieving these goals involves several significant hurdles related to the complex nature of the biology being studied, the models used to represent that biology, and the contextualization of those models in an approximation of a clinical environment. In this work, we address some of the challenges associated with the use of agent-based models (ABMs) to generate clinically-relevant simulation experiments, specifically those related to the effective and efficient parameterization of complex ABMs and the evaluation of those parameterizations within a clinical context that itself represents a multi-parametric space. We perform this work using a previously validated ABM of acute systemic inflammation (3, 4) while studying the clinical context of sepsis.

Sepsis is an inflammatory condition with a mortality rate of between 28%-50%(5). Numerous mechanistic computational simulations of acute inflammation and sepsis have been utilized over the past two decades(3, 6-12). These models have demonstrated that the sepsis population is much more heterogeneous than previously thought and this can be reflected by utilizing a range of multidimensional parameters that correlate to biologically plausible behaviors and phenotypes. Despite insights generated form these methods, there remain considerable challenges in the calibration and parameterization of the models. The description (contextualization) of the environment in which a biomedical simulation operates and parameterization of internal model rules (model content) requires the optimization of a large number of free-parameters; given the wide range of variable combinations, along with the intractability of ***ab initio*** modeling techniques which could be used to constrain these combinations, an astronomical number of simulations would be required to achieve this goal.

The primary model analyzed in this work is the Innate Immune Response Agent Based Model (IIRABM) (6, 13). The IIRABM is an abstract representation/simulation of the human inflammatory signaling network response to injury; the model has been calibrated such that it reproduces the general clinical trajectories seen in sepsis. The IIRABM operates by simulating multiple cell types and their interactions, including endothelial cells, macrophages, neutrophils, TH0, TH1, and TH2 cells as well as their associated precursor cells. The simulated system dies when total damage (defined as aggregate endothelial cell damage) exceeds 80%; this threshold represents the ability of current medical technologies to keep patients alive (i.e., through organ support machines) in conditions that previously would have been lethal. The IIRABM is initiated using 5 external variables – initial injury size, microbial invasiveness, microbial toxigenesis, environmental toxicity, and host resilience.

The model’s internal parameterization can be thought of as an abstraction of an *in vivo* genetic signaling network while the model context defines the simulated injury to which the model responds. This model has successfully replicated the range of cytokine time-course dynamics of sepsis and healing across a wide range of model contexts, however all models have been “genetically identical,” analogous to a typical mouse experiment. In practice, we recognize that there are two primary sources of variation in biological data. In any biological system, the response to a given stimulus contains some intrinsic degree of stochasticity – this can be seen in the range of responses from genetically identical mice to identical stimuli. Alternatively, variance in biological data output can arise from the genetic variability among individuals represented in the experiment. In order to determine the range of individuals/conditions our model can represent, internal parameterization boundaries (as well as their associated contextual boundaries) must be determined.

### Mapping as an alternative to sensitivity analysis

The problem of combinatorial complexity in the selection of model parameters is well-established in the computational/biological modeling communities (14-18). In previous work (3), we utilized high-performance computing to demonstrate the need for comprehensive “data coverage” among possible model states as well as the importance of internal parameter variation (as compared to model structure) to capture the full range of biological heterogeneity seen clinically. Here, we have extended that work to consider both model internal parameterizations as well as the parameterization which describes the model perturbation that generates disease. We posit that parameter space mapping, which has recently been rendered tractable with the rise of machine learning, has more generalizable utility than a traditional sensitivity analysis; the rationale for this statement is described below and in (19).

In a standard parameter sensitivity analysis (20-22), the dependence of mode output on variance in a single parameter (or potentially a combination or parameters) is quantified. In a complex ABM, this process requires the consideration of a large number of potentially free parameters, making a comprehensive calibration difficult (23-27) and significantly diminishing the utility of traditional parameter sensitivity analysis techniques (28, 29). These difficulties are compounded when considering the range of biological heterogeneity seen experimentally and clinically (3, 19).

Sensitivity analysis can be effective at characterizing certain dynamic processes of ABMs, however, when traditional sensitivity analysis is applied to the high-dimensional, complex parameter spaces seen in detailed ABMs, it precludes the consideration of alternate model rule configurations which can generate equally viable model outputs and behaviors. As an illustrative example, consider the following rule, present in the original version of the IIRABM, describing a term which contribute the differentiation switch from a Th0 cell into a Th1 or Th2 cell:

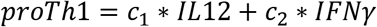

Where the *c_i_′s* represent some constant values used to weight the individual contributions of the array of cytokines. One could perform a sensitivity analysis on a model consisting solely of this rule, however that analysis would miss import bio-plausible model calibrations/parameterizations. While the rule is coded this way in the model, this representation does not actually represent the assertion made by the rule. A more complete way to write this rule would be:

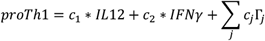

Where *j* sums over the complete set of cytokines on the model, Γ_*j*_ represents the concentration of a specific cytokine, and the constant weighting terms, *c_j_* = 0. Thus, a comprehensive sensitivity analysis must consider implicit zeros in model rule construction, which can vastly increase the size of the task. Additionally, the model may only be sensitive to certain parameters in a specific context. Consider a more generic version of the above rule (model parameterization 1), which hypothetically leads to biologically plausible model output:

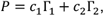

Which more completely, would be written:

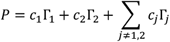

Where *c_j_* = 0. There is no supposition that this hypothetically calibrated rule is uniquely correct. Assume the existence of an additional/alternative calibrated rule (model parameterization 2), which leads to equally biologically plausible model output:

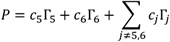

In a model parameterization 1, in which all cytokine multipliers are set to 0 except for cytokine 1 and cytokine 2, the model will appear more sensitive to cytokines 1 and 2 than the others. In model parameterization 2, the model will appear more sensitive to cytokine 5 and cytokine 6. A sensitivity analysis on one model parameterization is not valid for all other model parameterizations. A comprehensive sensitivity analysis would then have to incorporate the model configuration under which it is being performed, which is computationally intractable for all but the smallest of models. In order to render this task computationally tractable, we have employed a nested active learning approach in order to efficiently and comprehensively explore model parameter space.

## Methods

### EMEWS

During the initial code development, the AL models were trained and integrated using the Extreme-scale Model Exploration With Swift (EMEWS) framework (30-33). EMEWS enables the creation of high-performance computing (HPC) workflows for implementing large-scale model exploration studies. Built on the general-purpose parallel scripting language Swift/T (34), multi-language tasks can be combined and run on the largest open science HPC resources (35) via both data-flow semantics and stateful resident tasks. The ability that EMEWS provides for incorporating model exploration algorithms such as AL, implemented in R or Python, allows for the direct use of the many libraries relevant to ML that are being actively developed and implemented as free and open source software.

### Active Learning

Active learning (AL) is a sub-field of machine learning (ML) which focusses on finding the optimal selection of training data to be used to train a ML or statistical model (36). AL can be used for classification (37, 38) or regression (39, 40). AL is an ideal technique for modeling problems in which there is a large amount of unlabeled data and manually labelling that data is expensive. In these circumstances (specifically the costly data labelling) AL provides to most generalizable and accurate model for the cheapest cost, which for the purposes of this work, is computation time.

The lower-level AL procedure seeks to find the boundary of the parameter space deemed “clinically relevant” (3) as a function of four parameters which describe the context in which the IIRABM operates: two measures of microbial virulence (invasiveness and toxigenesis), host resilience, and environmental toxicity. In this scheme, there are two classes: clinically relevant and not clinically relevant. We assume that there exists some function,

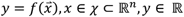

which accurately classifies model context parameters can be approximated given input data from the training set:

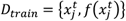

for *j* = 1,…,*n*. The NN model uses a binary cross-entropy (41) loss function, in which the loss is given by:

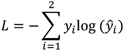

Where *y_i_* is the ground truth value and *ŷ_i_* is the NN-approximated score. The AL algorithm begins with a randomly selected set of 20 points. The IIRABM simulation then runs a fixed number of stochastic replicates of the input points to determine class membership. This information is then used to train the ML model. The algorithm then ranks the remaining unlabeled parameterizations by class-membership uncertainty (see Eq. 1).

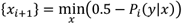

Those parameterizations whose class is most uncertain in the current ML model are then selected for labeling and the process repeats until a stopping criterion is reached; for the purposes of this work, once the cross-validation accuracy crossed 0.95, the algorithm was stopped.

The upper-level AL workflow uses a modified version of Dropout-based AL for regression presented in (40). The goal of this AL-workflow is twofold: to predict the volume of CR space and to predict the centroid location of CR-space, given a model internal parameterization. For each regression task, we assume that there exists a function,

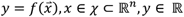

which approximates a map of CR space as a function of internal model parameterization which comprises the training set:

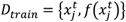

for *j* = 1,…,*n*. The NN model uses a mean-squared error (MSE) loss function, given by:

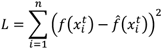

Where *f_i_* is the ground truth value for either the CR volume or centroid and 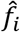 is value regressed by the NN. In this scheme, we utilize a four-layer fully-connected neural network with a 256-Dropout-128-Dropout architecture. The dropout layer (42) serves to provide a stochastic variability to the output of the NN.

**Figure 1:**
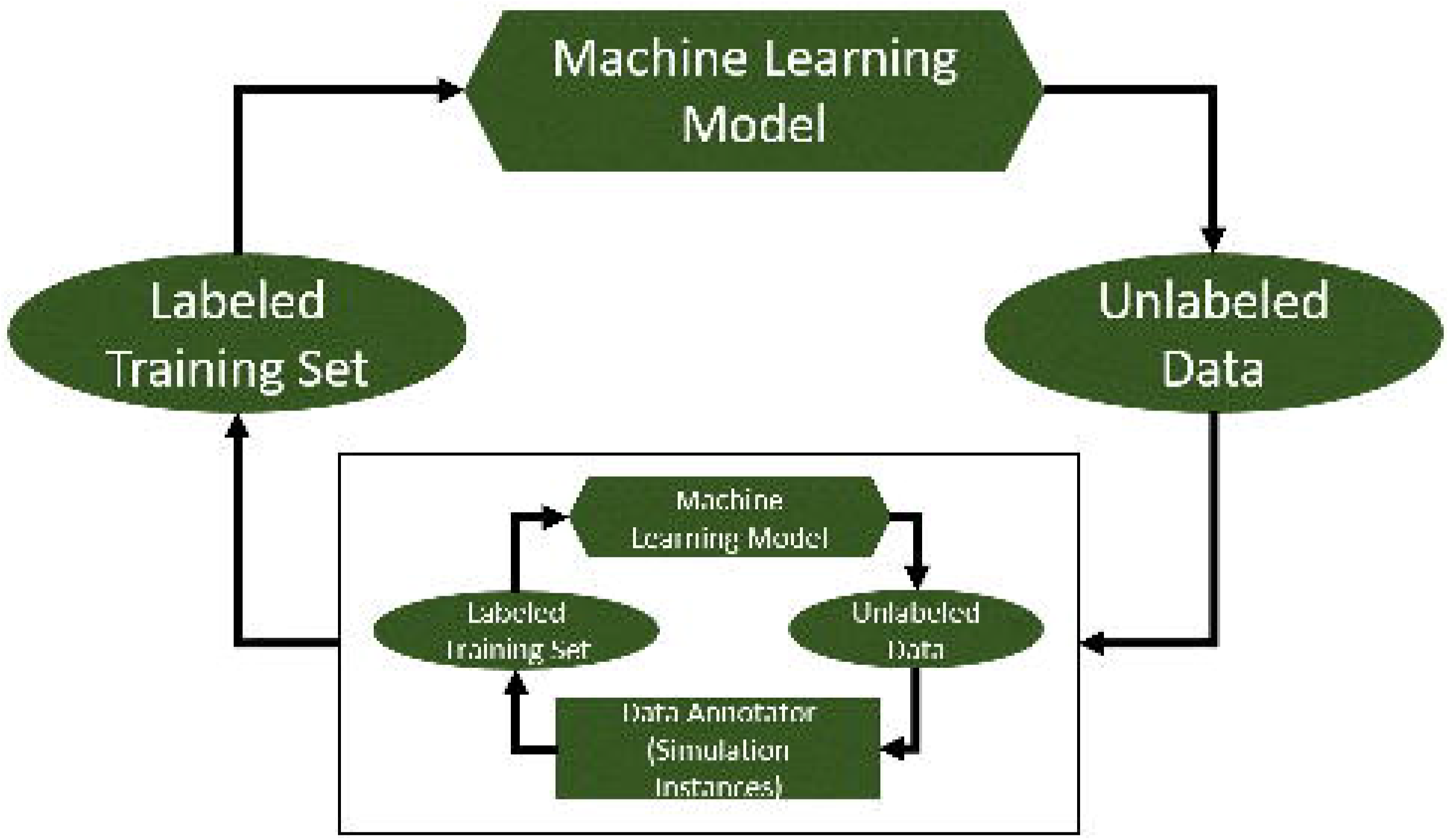
A diagram illustrating the nested active learning workflow

We discretize the model’s internal parameter space into 9 bins, representing augmentations or inhibitions to specific cytokine pathways, giving 40,353,607 potential internal model parameterizations. We begin by pre-selecting 10,000 of these internal parameterizations randomly; this random selection then makes up the available pool, 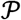, of unlabeled data. From this pool, we begin the AL procedure by selecting 100 internal parameterizations randomly from 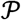. These internal parameterizations are then fed into the lower-level AL workflow, which is used to map the CR space and return an approximate volume and center-point. This data is then used to train the upper-level neural net (see Fig. 1). The trained NN is then used to predict the volume or centroid location for the remaining unlabeled data for 10 stochastic replicates (the dropout layer provides stochasticity). The parameterizations from 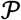 which have the highest variance are selected for labeling, and this process repeats. Pseudocode for this procedure is given below:

1. Initialize training pool 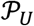; upper-level dataset *D_IP_*; *z_u_*, the maximum size of the dataset; and *m_u_* samples to be added on each iteration,
2. While |*D_IP_*| < *z_u_*:

a. For each element *i* in *D_IP_*:

i. Initialize training pool 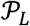; lower-level dataset *D_EP_*; *z_l_*, the maximum size of the dataset; and samples to be added on each iteration,
ii. While |*D_EP_*| < *z_l_*:

1. Train network on *D_EP_*
2. Obtain rank *r_j_* for each *x_j_* in 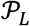 according to maximum class-uncertainty
3. Label the set of *m_l_* parameterizations from 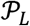
4. Add the annotated data to *D_EP_*
5. Calculate stopping metrics, stop if appropriate
b. Train network on *D_IP_*
c. Obtain rank *r_i_* for each *x_i_* in 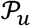 according to maximum regression variance
d. Label the set of *m_u_* parameterizations from 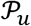
e. Add the annotated data to *D_IP_*
f. Calculate stopping metrics, stop if appropriate

Source code and input data can be found at: https://github.com/An-Cockrell/IIRABM_MRM_AL/

## Results

In the lower-level AL workflow, we map CR space as a function of four parameters, external to the IIRABM’s internal rule set. An example of this space can be seen in Fig. 2. In this figure, outcome spaces for patients with low environmental toxicity (toxicity=1) to high environmental toxicity (toxicity=10) are shown from left to right. Each point represents 4000 *in silico* patients (40 injury sizes, 100 stochastic replicates). Points are color-coded based on the outcomes generated. The CR space is shown in green.

**Figure 2:**
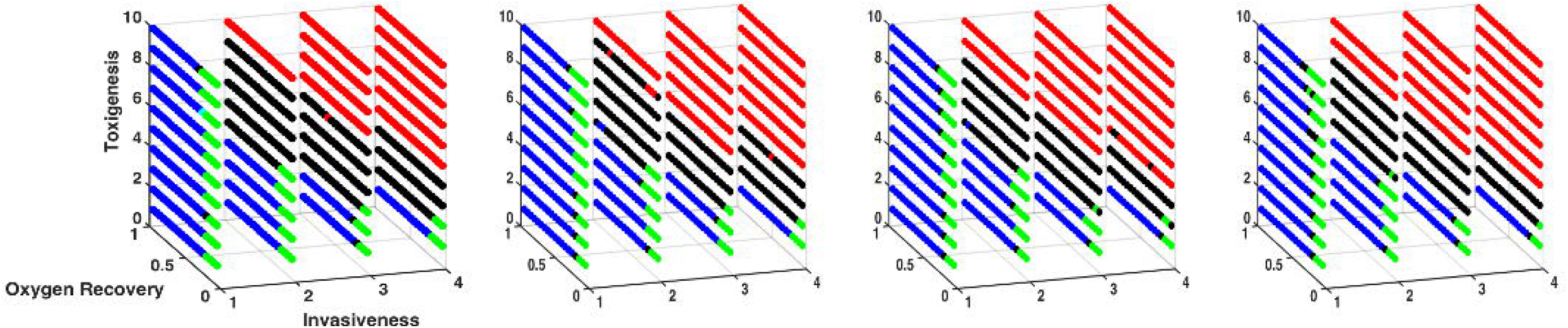
Outcome spaces for patients with low environmental toxicity (toxicity=1) to high environmental toxicity (toxicity=10) are shown from left to right. Each point represents 4000 *in silico* patients (40 injury sizes, each with 100 stochastic replicates). Points are color-coded based on the outcomes generated. Blue points represent simulations that healed under all circumstances. These points are distributed in regions of space where host resilience is high and the bacterial virulence is low (lower invasiveness and lower toxigenesis). Red points represent simulations that always died from overwhelming infection; these points are distributed in regions of high bacterial virulence (higher values for invasiveness and toxigenesis). Black points represent simulations that either died from overwhelming infection or healed completely and mark the boundary between simulations that always heal and simulations always die from infection. Pink points represent simulations which either died from overwhelming infection or hyperinflammatory system failure; these points are found primarily in the simulations that were treated with antibiotics and had low values for environmental toxicity and host resilience. Green points represent the Clinically Relevant simulations as these parameter sets lead to all possible outcomes; these points are distributed in regions of low to middle values of the host resilience parameter and moderately virulent infections. For all classes of simulation, the final outcomes are primarily dependent on the host resilience and microbial virulence

We utilized seven different ML models to map the CR space: Artificial Neural Network (43), Adaptive Boosting (44), Naïve Bayesian (45), Random Forest (46), TreeBag (47), AdaBoost M1 (48), Bag – Flexible Discriminant Analysis with Generalized Cross Validation (49). We compare an ensemble of models to ensure we have selected an appropriate ML strategy to explore the model in the most efficient manner. Results from this are shown in Fig. 3, which displays the F-score as a function of AL iteration number (and by proxy, dataset size).

**Figure 3:**
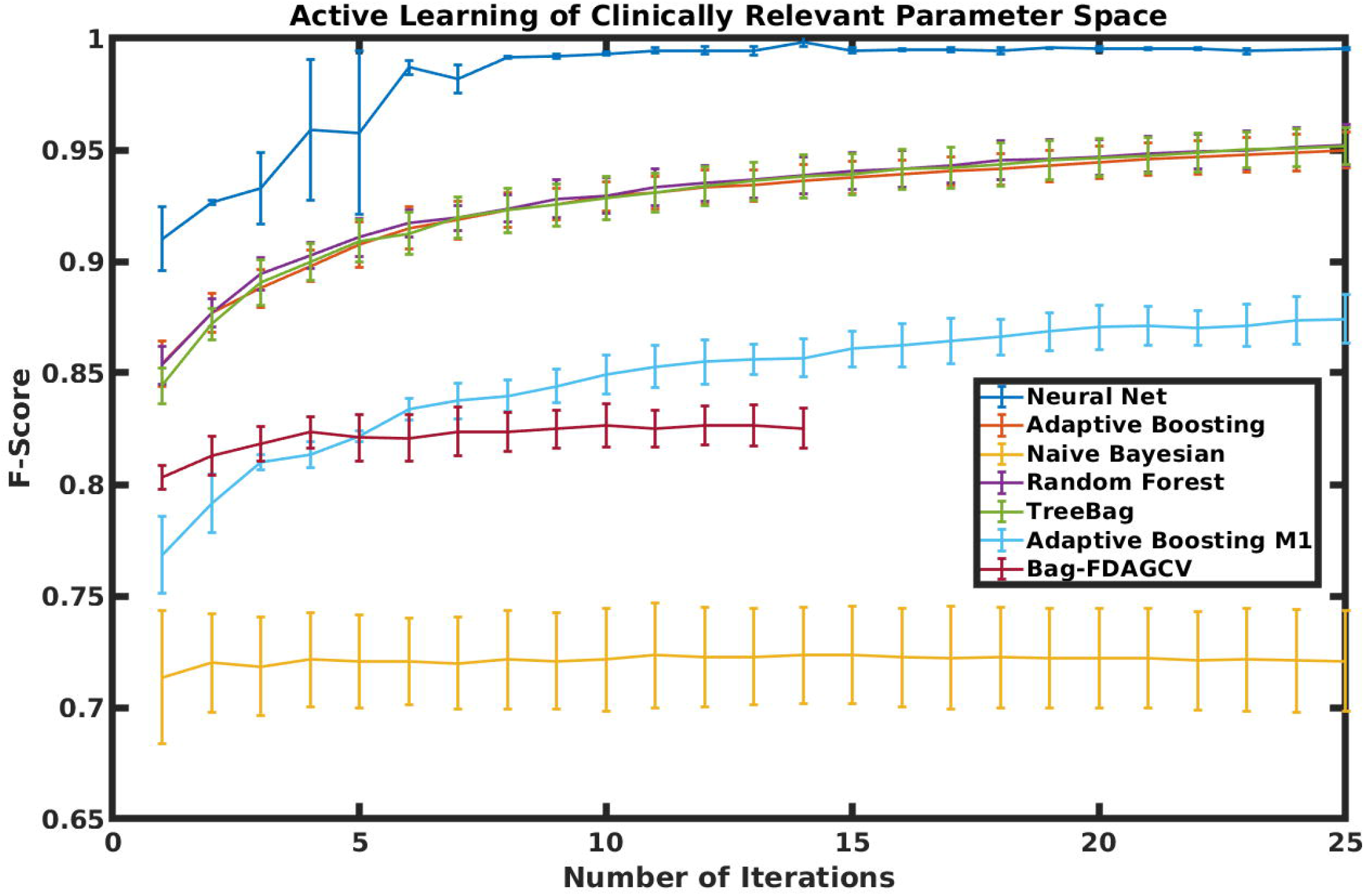
Results from Lower-Level AL. Clinically Relevant space (see Fig. 2) is mapped as a function of four parameters, external to the IIRABM’s internal rule set. We utilized seven different ML models to map the CR space: Artificial Neural Net, Adaptive Boosting, Naïve Bayesian, Random Forest, TreeBag, AdaBoost M1, Bag – Flexible Discriminant Analysis with Generalized Cross Validation. The F-score is shown on the y-axis as a function of the number of AL iterations performed.

It is readily apparent that a NN is the best type of ML model for mapping this space. By iteration 10, which uses 1000 parameterizations (out of 8800 possible), we can achieve an average class-prediction accuracy of >98%. The resulting ML model is then utilized to efficiently calculate the location and centroid of the CR space and train the upper-level neural net. This is due to the ease with which NN’s can approximate nonlinear functions (50). We present an illustration of this in Figure 4, which displays three unique clinically relevant spaces for three unique internal model parameterizations, in a threedimensional slice of a four-dimensional perturbation-parameter space. The legend refers to an exponent determining the strength of augmentation or inhibition for all pathways; the meaning of and rationale for this exponent are described in detail in (6). Red points labelled as ‘Unique −1’ are those points that are unique to the parameterization in which all protein synthesis pathways are inhibited by 90% (10^−1^ of the uninhibited pathway secretion). Green points are unique to the parameterization in which all protein synthesis pathways are augmented by a factor of 10^1^. Teal points are unique to the parameterization in which all protein synthesis pathways are unchanged. Black points are those shared by the maximally inhibited and unchanged protein synthesis pathways.

**Figure 4:**
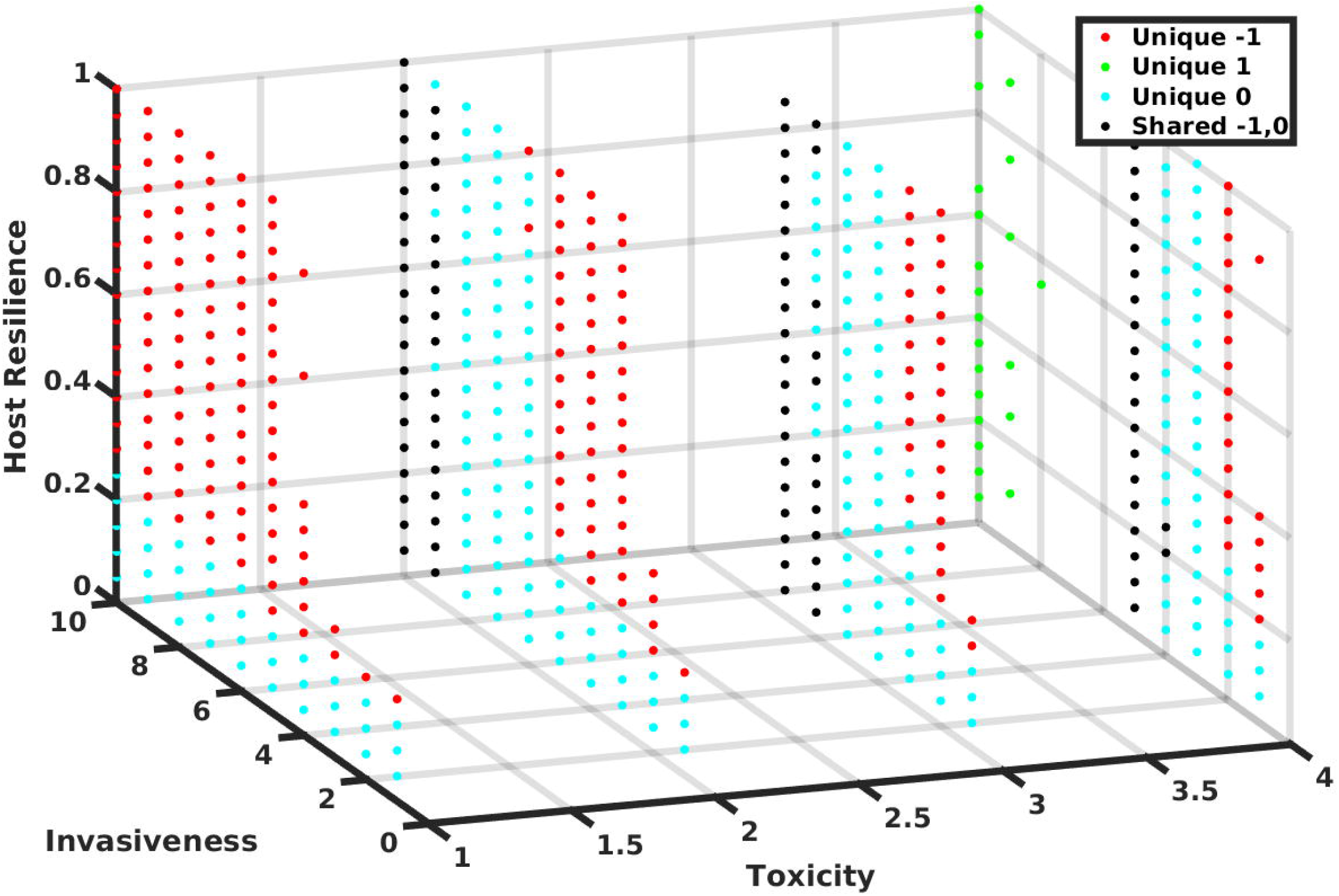
Clinically Relevant Space for Varying Internal Parameterizations: Clinically Relevant space for 3 unique internal parameterizations is mapped as a function of three parameters external to the IIRABM’s rule-set, with the environmental toxicity parameter held fixed at 2 (a low level). The legend refers to an exponent determining the strength of augmentation or inhibition for all pathways. Red points labelled as ‘Unique −1’ are those points that are unique to the parameterization in which all protein synthesis pathways are inhibited by 90% (10^−1^ of the uninhibited pathway secretion). Green points are unique to the parameterization in which all protein synthesis pathways are augmented by a factor of 10^1^. Teal points are unique to the parameterization in which all protein synthesis pathways are unchanged. Black points are those shared by the maximally inhibited and unchanged protein synthesis pathways.

Results from the upper-level AL procedure are shown in Figure 5. In panel A, we display the percent volume error as a function of the number of training samples for AL, Random Sampling (RS), and “Actively Not Learning” (ANL). ANL refers to utilizing the opposite of the AL sampling criterion. In this case, for AL we chose samples that maximized prediction variance; for ANL, we chose samples that minimized prediction variance. As expected, AL outperforms RS and requires fewer samples to converge to the error minimum. Additionally, both methods significantly outperform ANL, as expected. In Panel B, we show the standard deviation of the error for the previous three methods. Here, AL significantly outperforms RS in that the intelligent sampling criterion leads to a suite of models with a larger degree of precision in the volume prediction, whereas the changes in standard deviation of the error are minimal for ANL and minimal for RS after the first few samples. Panel C displays the error (as a Euclidean distance from the predicted centroid point to the true value) as a function of the number of samples. Once again, AL outperforms RS, though by a relatively modest amount.

**Figure 5:**
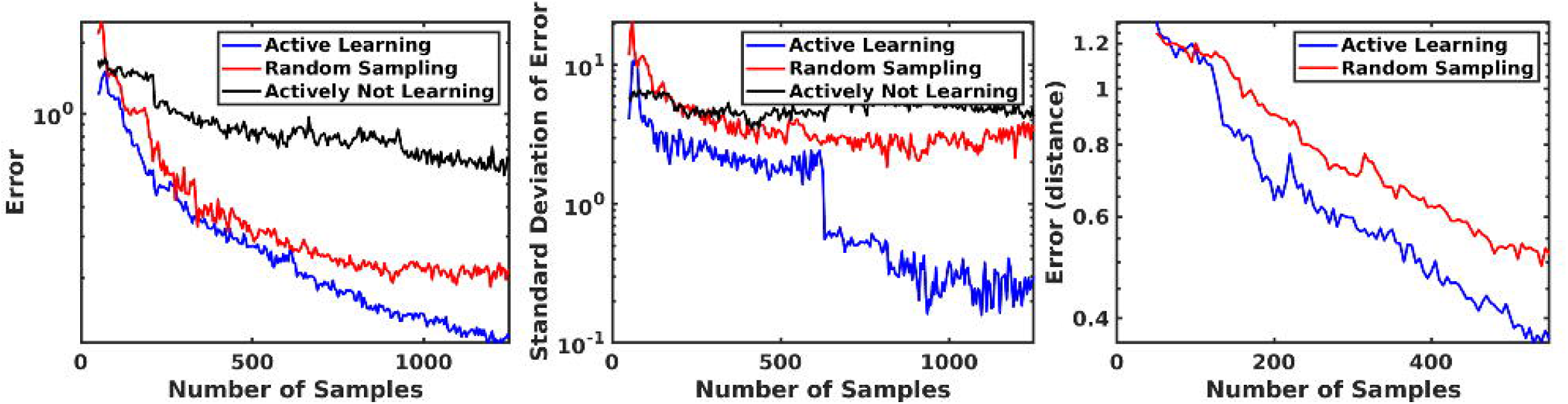
Results from Upper-Level AL. In Panel A, we show the percent volume error as a function of the number of input training samples using Active Learning (AL), Random Sampling (RS), and Actively Not Learning (ANL), in which the learning criterion is the opposite of the AL criterion. We see that AL arrives at a more accurate prediction with fewer samples than RS or ANL. In Panel B, we show the standard deviation of the error of the volume prediction for the three above methods and note that AL not only generates a suite of more accurate models, but also has a much higher degree of precision. In Panel C, we show the error (in this case a Euclidean distance) in the centroid location prediction. AL once again outperforms RL.

## Discussion

One of the primary goals and benefits of agent-based modeling is to use the computational model as an experimental proxy system (51) detailed enough to capture the vital aspects of patients and their associated clinical setting. The quest for increasingly detailed representation when using ABMs is a manifestation of the concurrent increase in mechanistic knowledge acquired from ongoing research; the consequence of these highly detailed models is that they contain a multitude of parameters within a highly connected set of rules and/or equations. The perception of intractability of effectively parameterizing such models is considered a major limitation in their use. Our prior work has demonstrated the need to operate with these models across a wide range of their parameter space, which we consider a representation of genetic variability with regards to pathway responsiveness among a clinical population (3) the work presented herein provides a demonstration of methods with which these parameter spaces can be identified.

We have described a nested active learning workflow which efficiently and accurately can characterize a high-dimensional and complex mechanism-based multi-scale model. We remove inefficiencies due to oversampling small regions of parameter space using the Monte-Carlo Dropout Uncertainty Estimation approach (40). We note that AL outperforms RS in both the volume and centroid location predictions, but the greatest strength comes from the significant increase in precision generated by a suite of AL-trained models. This work demonstrates that comprehensive (and accurate) exploration of computational models with many parameters is both possible and computationally tractable, given current techniques in machine learning and high-performance computing. Additionally, while in some circumstances, the gains offered by AL are modest, it does help to minimize the cost of the computation.

In order to increase the utility of this work, we will develop higher resolution maps by labeling external parameter points individually rather than regressing the volume and centroid of the CR space. This will likely include collapsing the nested AL structure into a single network which takes internal and external parameterizations simultaneously, sacrificing some efficiency gained by the nested structure for increased precision in class identification. Pre-trained models can be incorporated into the development and training of control strategies by enabling the selection of disease conditions with specific cytokine dynamics and mortality rates without a computationally expensive search process.

The ultimate goal of this method when applied to any computational model would be the generation of a relatively compact data structure, in this case, one or more pre-trained neural networks, which accurately predicts some feature of interest to the computational model; in this case, that feature is whether or not a specific individual (internal model parameterization/model content) will certainly live or die when it experiences a specific perturbation (perturbation parameterization/model context). The appropriate threshold for ‘accuracy’ can be determined based on the specific requirements of the application to which it is applied as well as considering the diminishing returns to ML-model prediction accuracy as the number of training samples increases.

## Acknowledgements

This research is based upon work supported by the U.S. Department of Energy, Office of Science, under contract DE-AC02-06CH11357 and the National Institutes of Health grant U01EB025825. This project used high performance computing resources of the National Energy Research Scientific Computing Center, a DOE Office of Science User Facility supported by the Office of Science of the U.S. Department of Energy under Contract No. DE-AC02-05CH11231, as well as resources provided by the Vermont Advanced Computing Core (VACC). Additionally, this research was supported in part by NIH through resources provided by the University of Chicago Computation Institute (Beagle2) and the Biological Sciences Division of the University of Chicago and Argonne National Laboratory, under grant 1S10OD018495-01 and under contract DE-AC02-06CH11357.

## References

1. Gambhir SS, Ge TJ, Vermesh O, Spitler R. Toward achieving precision health. Sci Transl Med. 2018;10(430):eaao3612.

2. Morrison TM, Pathmanathan P, Adwan M, Margerrison E. Advancing regulatory science with computational modeling for medical devices at the FDA’s Office of Science and Engineering Laboratories. Frontiers in medicine. 2018;5.

3. Cockrell C, An G. Sepsis reconsidered: Identifying novel metrics for behavioral landscape characterization with a high-performance computing implementation of an agent-based model. J Theor Biol. 2017;430:157–68.

4. An G. In silico experiments of existing and hypothetical cytokine-directed clinical trials using agent-based modeling. Critical care medicine. 2004;32(10):2050–60.

5. Wood KA, Angus DC. Pharmacoeconomic implications of new therapies in sepsis. Pharmacoeconomics. 2004;22(14):895–906.

6. Cockrell RC, An G. Examining the controllability of sepsis using genetic algorithms on an agent-based model of systemic inflammation. PLoS Comput Biol. 2018;14(2):e1005876.

7. An G, Nieman G, Vodovotz Y. Computational and systems biology in trauma and sepsis: current state and future perspectives. Int J Burns Trauma. 2012;2(1):1–10.

8. Goldman D, Bateman RM, Ellis CG. Effect of sepsis on skeletal muscle oxygen consumption and tissue oxygenation: interpreting capillary oxygen transport data using a mathematical model. American Journal of Physiology-Heart and Circulatory Physiology. 2004.

9. Vodovotz Y, Billiar TR. In Silico Modeling: Methods and Applications toTrauma and Sepsis. Critical care medicine. 2013;41(8):2008.

10. Vodovotz Y. Computational modelling of the inflammatory response in trauma, sepsis and wound healing: implications for modelling resilience. Interface Focus. 2014;4(5):20140004.

11. Siqueira-Batista R, Gomes A, Possi M, Oliveira A, Sousa F, Silva C, et al., editors. Computational modeling of sepsis: perspectives for in silico investigation of antimicrobial therapy. II International Conference on Antimicrobial Research-ICAR2012 Lisbon (Portugal); 2012.

12. An G, Nieman G, Vodovotz Y. Toward computational identification of multiscale “tipping points” in acute inflammation and multiple organ failure. Annals of biomedical engineering. 2012;40(11):2414–24.

13. Cockrell C, Christley S, An G. Investigation of inflammation and tissue patterning in the gut using a Spatially Explicit General-purpose Model of Enteric Tissue (SEGMEnT). PLoS Comput Biol. 2014;10(3):e1003507.

14. Karp RM. On the computational complexity of combinatorial problems. Networks. 1975;5(1):45–68.

15. Sneddon MW, Faeder JR, Emonet T. Efficient modeling, simulation and coarse-graining of biological complexity with NFsim. Nat Methods. 2011;8(2):177–83.

16. Hopfield JJ, Tank DW. “Neural” computation of decisions in optimization problems. Biological cybernetics. 1985;52(3):141–52.

17. Edwards R, Glass L. Combinatorial explosion in model gene networks. Chaos. 2000;10(3):691–704.

18. Neumann F, Witt C. Combinatorial optimization and computational complexity. Bioinspired Computation in Combinatorial Optimization: Springer; 2010. p. 9–19.

19. Cockrell C, An G. Genetic Algorithms for model refinement and rule discovery in a highdimensional agent-based model of inflammation. bioRxiv. 2019:790394.

20. Saltelli A, Ratto M, Andres T, Campolongo F, Cariboni J, Gatelli D, et al. Global sensitivity analysis: the primer: John Wiley & Sons; 2008.

21. Cukier R, Levine H, Shuler K. Nonlinear sensitivity analysis of multiparameter model systems. Journal of computational physics. 1978;26(1):1–42.

22. Saltelli A, Tarantola S, Campolongo F, Ratto M. Sensitivity analysis in practice: a guide to assessing scientific models. Chichester, England. 2004.

23. Ling Y, Mahadevan S. Quantitative model validation techniques: New insights. Reliability Engineering & System Safety. 2013;111:217–31.

24. Macal CM, editor Model verification and validation. Workshop on” Threat Anticipation: Social Science Methods and Models; 2005.

25. Calvez B, Hutzler G, editors. Parameter space exploration of agent-based models. International Conference on Knowledge-Based and Intelligent Information and Engineering Systems; 2005: Springer.

26. Abramson D, Bethwaite B, Enticott C, Garic S, Peachey T, editors. Parameter space exploration using scientific workflows. International Conference on Computational Science; 2009: Springer.

27. Carley KM. Validating computational models. Paper available at http://www.casos.cs.cmu.edu/publications/papers.php. 1996.

28. Saltelli A, Annoni P. How to avoid a perfunctory sensitivity analysis. Environmental Modelling & Software. 2010;25(12):1508–17.

29. Ratto M, Pagano A, Young P. State dependent parameter metamodelling and sensitivity analysis. Computer Physics Communications. 2007;177(11):863–76.

30. Ozik J, Collier N, Heiland R, An G, Macklin P. Learning-accelerated discovery of immune-tumour interactions. Mol Syst Des Eng. 2019;4(4):747–60.

31. Ozik J, Collier NT, Wozniak JM, Macal C, An G. Extreme-scale Dynamic Exploration of a Distributed Agent-based Model with the EMEWS Framework. IEEE Trans Comput Soc Syst. 2018;5(3):884–95.

32. Ozik J, Collier N, Wozniak JM, Macal C, Cockrell C, Friedman SH, et al. High-throughput cancer hypothesis testing with an integrated PhysiCell-EMEWS workflow. BMC Bioinformatics. 2018;19(Suppl 18):483.

33. Ozik J, Collier NT, Wozniak JM, Spagnuolo C, editors. From desktop to large-scale model exploration with Swift/T. 2016 Winter Simulation Conference (WSC); 2016: IEEE.

34. Wozniak JM, Armstrong TG, Wilde M, Katz DS, Lusk E, Foster IT, editors. Swift/t: Large-scale application composition via distributed-memory dataflow processing. 2013 13th IEEE/ACM International Symposium on Cluster, Cloud, and Grid Computing; 2013: IEEE.

35. Wozniak JM, Jain R, Balaprakash P, Ozik J, Collier NT, Bauer J, et al. CANDLE/Supervisor: a workflow framework for machine learning applied to cancer research. BMC Bioinformatics. 2018;19(Suppl 18):491.

36. Cohn DA, Ghahramani Z, Jordan MI. Active learning with statistical models. Journal of artificial intelligence research. 1996;4:129–45.

37. Brinker K. On active learning in multi-label classification. From Data and Information Analysis to Knowledge Engineering: Springer; 2006. p. 206–13.

38. Huang S-J, Jin R, Zhou Z-H, editors. Active learning by querying informative and representative examples. Advances in neural information processing systems; 2010.

39. Schein AI, Ungar LH. Active learning for logistic regression: an evaluation. Machine Learning. 2007;68(3):235–65.

40. Tsymbalov E, Panov M, Shapeev A, editors. Dropout-Based Active Learning for Regression. International Conference on Analysis of Images, Social Networks and Texts; 2018: Springer.

41. De Boer P-T, Kroese DP, Mannor S, Rubinstein RY. A tutorial on the cross-entropy method. Annals of operations research. 2005;134(1):19–67.

42. Srivastava N, Hinton G, Krizhevsky A, Sutskever I, Salakhutdinov R. Dropout: a simple way to prevent neural networks from overfitting. The Journal of Machine Learning Research. 2014;15(1):1929–58.

43. White H. Artificial neural networks: approximation and learning theory: Blackwell Publishers, Inc.; 1992.

44. Rojas R. AdaBoost and the super bowl of classifiers a tutorial introduction to adaptive boosting. Freie University, Berlin, Tech Rep. 2009.

45. Kononenko I, editor Semi-naive Bayesian classifier. European Working Session on Learning; 1991: Springer.

46. Ho TK, editor Random decision forests. Proceedings of 3rd international conference on document analysis and recognition; 1995: IEEE.

47. Breiman L. Bagging predictors. Machine learning. 1996;24(2):123–40.

48. Freund Y, Schapire RE, editors. Experiments with a new boosting algorithm. icml; 1996: Citeseer.

49. Hastie T, Tibshirani R, Buja A. Flexible discriminant analysis by optimal scoring. Journal of the American statistical association. 1994;89(428):1255–70.

50. Barron AR. Universal approximation bounds for superpositions of a sigmoidal function. IEEE Transactions on Information theory. 1993;39(3):930–45.

51. An G, Fitzpatrick BG, Christley S, Federico P, Kanarek A, Neilan RM, et al. Optimization and Control of Agent-Based Models in Biology: A Perspective. Bull Math Biol. 2017;79(1):63–87.

